# Characterization of a novel gene-environment-based animal model to study resilience and susceptibility to PTSD and co-morbid depression

**DOI:** 10.1101/2022.09.14.507883

**Authors:** Lia Parada Iglesias, Arthur Alves Coelho, Nicole Rodrigues da Silva, Heidi K. Müller, Fabricio A. Moreira, Gregers Wegener, Sâmia Joca

**Affiliations:** Translational Neuropsychiatry Unit, Department of Clinical Medicine, Aarhus University; Ribeirao Preto Medical School, University of Sao Paulo; Department of Biomedicine, Aarhus University; Pharmacology Department, Universidade Federal de Minas Gerais

**Keywords:** Flinders Sensitive/Resistant Line Rats, hippocampus, fear, memory, depression, PTSD, FKBP51, corticosterone

## Abstract

**BACKGROUND:** Post-traumatic stress disorder (PTSD) and co-morbid depression are frequently associated with severe symptoms, poor response to treatment and worse prognosis. Due to the absence of a suitable animal model, little is known about the biological basis of the comorbidity, severely limiting the discovery of new and more effective treatment options. The Flinders Sensitive Line rats (FSL) is a well-validated, selectively bred animal model of depression. However, several of its features, such as cognitive deficits and altered hypothalamic-pituitary-adrenal (HPA) axis response, also match symptomatic clusters of PTSD. In parallel, its resistant counterpart, the Flinders Resistant Line (FRL), is extensively used as a simple control. Still, little is known about its performance compared to the original strain, Sprague Dawley (SD), from which the FSL/FRL was originally derived.

**AIMS:** Characterizing the behavioural performance and mechanisms involved in FSL, FRL and SD rats in fear-memory paradigms.

**METHODS:** FSL, SD and FRL animals were submitted to tests assessing hippocampal-dependent and fear-related memory. Subsequently, plasticity factors and endocrine responses to stress were analysed to elucidate the molecular basis for the observed behavioural alterations.

**RESULTS:** We found that FRL animals presented intact recognition memory and innate fear responses but could not properly display conditioned responses in the Conditioned Fear Conditioning (CFC) paradigm. FSL animals, despite a poor performance in the Novel Object Recognition task (NOR), showed similar levels of conditioned responses compared to SD, but impairments in extinction learning, a feature highly related to PTSD. The behavioural alterations were accompanied by alterations in plasma corticosterone levels and hippocampal expression of the glucocorticoid receptor and FKBP51.

**CONCLUSION:** For the first time, we demonstrate an animal model of resilience and vulnerability to PTSD and co-morbid depression. The results suggest that the endophenotypes may be based on aberrant endocrine stress responses in the hippocampus.

## 1 INTRODUCTION

On average, 70% of people will experience a traumatic event at some point in their life (1, 2). Although most people will fully recover after the initial stress response, 3.9% will develop an aberrant representation of aversive memory constructs characterised by over-consolidation, memory generalisation and impaired extinction, resulting in persistent reminding of the traumatic experience and emotional consequences characteristics of post-traumatic stress disorder (PTSD) (3). PTSD patients have a higher risk of developing other psychiatric disorders, among which Major Depressive Disorder (MDD) (4–7) is the most prevalent, with 50-70% of the patients presenting this comorbidity (4–6). PTSD and co-morbid depression is a severe clinical condition characterised by severe symptoms, poor response to treatment and worse prognosis (8–10). It is, however, unclear whether MDD is a consequence of experiencing PTSD or an independent risk factor for developing PTSD following trauma exposure (7). Unfortunately, the neurobiological basis of PTSD-MDD comorbidity remains poorly understood, partly due to the lack of appropriate animal models, which hinders the development of better treatment options.

Both PTSD and MDD patients present mal-adaptative expression of fear memory, cognitive deficits and increased negative bias (11–15),suggesting a common neurobiological substrate. The hippocampus (HPC) is part of a brain network involved in memory processing and has a central role in terminating the endocrine response to stress and promoting behavioural adaptation (16, 17). Interestingly, both PTSD and MDD patients present hippocampal dysfunctions, characterised by atrophy and synaptic loss (18–20), impaired neurogenesis (14, 21, 22), and impaired synaptic plasticity (23–25). A recent metanalysis of human studies suggested that the neuroplastic changes in the HPC are associated with impaired hippocampal-dependent tasks and constitute a vulnerability factor associated with PTSD and MDD development (26). Accordingly, a malfunctioning HPC has also been linked to the neuroendocrine dysregulations associated with PTSD and MDD (blunted or exacerbated HPA axis reactivity, respectively), which are also associated with increased risk of disease development (27, 28). Therefore, the trauma is necessary but not sufficient to trigger PTSD development (29); instead, the vulnerability pre-trauma, determined by cognitive and neurobiological factors, will moderate the susceptibility of some individuals to developing the disorder.

Although the currently available animal models of PTSD have been valuable tools for understanding the mechanisms of aversive memory processing and fear conditioning, they, unfortunately, have important translational limitations compromising the successful development of more effective treatments (29, 30). For instance, the absence of resilient subgroups and the lack of comorbidities, which are “more a rule than an exception” for MDD, are crucial limitations (29, 30). Consequently, studies of responses to traumatic events in animals that display proper endophenotypic characteristics of resilience or susceptibility to stress may provide valuable information about the neurobiology of PTSD and PTSD-MDD comorbidity.

The FSL, and its counterpart, the Flinders Resistant Line (FRL) rats, are inbred lines obtained by selective breeding from Sprague-Dawley (SD) rats based on their response to anticholinesterase agents (31). Years of research have demonstrated that the FSL display increased vulnerability to stress and depressive-like behaviours when compared to their control line, the FRL, and is considered a valid model to study gene x environment interactions in depression (31). For instance, FSL showed increased immobility in the forced swimming test, anhedonia (32, 33), increased rapid-eye-movement (REM) sleep (34, 35), cognitive deficits (36–41) and altered hippocampal structure (42). Some of these features are also common in PTSD, such as amygdala hyperreactivity against threats (43) or resistance to treatment after stress (44). Furthermore, chronic treatment with selective serotonin reuptake inhibitors (SSRIs), the first-line treatment for both PTSD and MDD, attenuates depressive-like behaviour (31) and emotional memory impairments in FSL animals (45). Despite the evidence mentioned above, little is known regarding the performance of FSL animals in behavioural tests that assess traumatic memory. Despite noticeable differences, much less is known about the FRL, which is often used interchangeably with SD rats (46). Therefore, our overall aim was to investigate the behaviour of FSL, FRL and SD rats in the contextual fear conditioning (CFC) paradigm, as well as the potential molecular pathways involved in their performance, to characterize a model vulnerability/resilience to PTSD-MDD to establish a new paradigm to study PTSD and MDD comorbidity.

## 2 MATERIAL AND METHODS

### 2.1 Animals

Nine to eleven-weeks-old male rats were used in the experiments. The FRL and FSL rats were kept in the breeding colonies at the Translational Neuropsychiatric Unit (TNU), Aarhus University, Denmark, originally derived from the FSL/FRL colonies at the University of North Carolina at Chapel Hill, USA, and Karolinska Institute, Stockholm, Sweden. Sprague-Dawley breeding pairs were obtained from Taconic Denmark A/S and bred at TNU. The rats were housed in pairs in Eurostandard Type IIIH cages with raised lids, shelter and nesting material, and free access to food and tap water. The cages were kept in a temperature-controlled room (21±1°C) with a standard dark-light cycle of 12/12h (lights on 06:00), and all the experiments were performed during the light phase of the cycle. All the procedures performed were approved by the Danish National Committee for Animal Experimentation in accordance with the EU Directive 2010/63/EU and the Danish regulations for experiments with animals.

### 2.2 Behavioural Tasks

The animals were handled for three days before each experiment and were moved to the behavioural room at least 1h before the start of the experiment and returned to the housing room 30 min after the end of the experiment. The animals of each phenotype were randomly assigned to different behavioural conditions and testing times to avoid experimental biases.

#### 2.2.1 Contextual Fear Conditioning

The experiment was carried out using the multiconditioning system (Multi Conditioning System, Version 1.0, TSE Systems GmbH, Bad-Homburg, Germany). The box was kept at 21°C, the background noise was set at 40-50dB, and the light intensity on the box floor was 70lx. All phases were separated by 24h. The following paradigm was applied: Day 1: habituation in the box for 10 min. Day 2, fear conditioning: 3 minu of habituation followed by 3 footshocks (1mA, 1s each, 40-60s random interval). The animals were kept in the box for one more minute before being returned to the homecages. Day 3, test: the animals were re-exposed to the conditioning context for 10 min. Days 4, 5 and 6, extinction sessions: each extinction training lasted 20 min, with 24h intervals. After the last extinction session, day 7, a reinstatement session was performed, in which the animals were allowed to explore the context for 3 min, followed by one shock (1mA, 1s) and 1 min habituation before returning to their homecages. Day 8: reinstatement was tested with a 5 min context exposure. The TSE Systems multiconditioning software recorded and analysed the behaviour each day.

#### 2.2.2 Auditory Fear Conditioning

The experiment was carried out using the multiconditioning system (Multi Conditioning System, Version 1.0, TSE Systems GmbH, Bad-Homburg, Germany). The box was kept at 21°C, the background noise was set at 40-50dB, and the light intensity on the box floor was 70lx. All the phases were separated by a 24h period. The following paradigm was applied: Day 1: the animals were allowed to habituate to the square box for 10 min. Day 2, conditioning: 3 min of habituation followed by three tones (30s, 2.8 kHz, 85dB, 180 or 100 intervals) co-terminating with a 1 s footshock (1mA). After the last shock, the animal was kept in the box for one more minute. Day 3, tone test: the animals were located in a circular arena and, after 3 min, presented to the three tones separated apart 180 or 100s, the animal remained in the arena 1 min after the last tone. Day 4: the context test was performed where the animal was exposed to the square box for 10 min.

#### 2.2.3 Novel Object Recognition

The test was performed as previously described (47). The following paradigm was applied: Day 1 (habituation): the animals were allowed to explore the box for 10 min freely. Day 2 (training): the animals were placed again in the box, two identical objects were introduced, and the animals were allowed to explore the objects for 5 min. Day 3 (test): the animals were placed in the box with one familiar object (same as in training) and one novel object and were allowed to explore the objects for 5 min. Exploring was considered when: the rat was directly interacting with the object; or with the nose in the direction of the object (1cm); climbing the object or touching it to explore above it was not considered exploration time (47). Animals that did not explore 30s during the training or 10s during the test were excluded (48). A blinded experimenter manually recorded the time spent with the objects. The exploration was calculated as a percentage of time exploring the novel object during the test phase (47, 48).

### 2.3 Molecular and histology procedures

The animals were taken to a separate room and euthanized by decapitation, i) immediately after the reinstatement (for dendritic spine analysis and HPLC analysis), or ii) 30 min after the conditioning (for western blot and ELISA). Following extraction of the brain, the HPC was rapidly dissected and frozen on dry ice powder and further processed as described below.

#### 2.3.1 HPLC

The samples were kept at −80°C until further processing for HPLC. Sample processing: 0.2M of HClO4 was added to the sample (x5 times the weight-mg in uL), followed by sonication of the tissue and centrifuged (14000G, 4°C, 10 min). The supernatant was collected and centrifuged again (14000G, 4°C, 1 min) in a 0.2μl filter. The experiment was run in two ThermoScientific ultimate 3000 uHPLC systems, one based on electrochemical detection (monoamines), 3000RS detector, and the other based on a fluorescent detector (glutamate), FLD-3100. For glutamate analysis, the samples were diluted at 2% in HCIO4 0.2M. The mobile phases for glutamate quantification consisted of two mobile phases (A and B), with A being a 20mM Sodium phosphate (pH 7.2) and B a 1:1 acetonitrile and methanol. The start mixture was 97% A and 3% B, with increasing amounts of B throughout the run with a slope of 5. The mobile phase for monoamines quantification was isocratic and consisted of sodium dihydrogen phosphate 0.08M, 1-Octanesulfonic acid sodium salt 2mM, acetonitrile 10%, EDTA 0.02mM and triethylamine 0.01%, the pH of the final solution was adjusted (pH 3). uHPLC analyzed samples with a column Kinetex 2,6μm EVO C18 100Å, Size LC Column 150 x 4,6 mm, (Phenomenex USA).

#### 2.3.2 Golgi-Cox staining

The procedure was performed using the FD Rapid GolgiStain kit (FD Neurotechnologies INC). One hemisphere was collected immediately after the reinstatement session, rinsed with Milli-Q water and immersed in the impregnation solution (solutions A and B, 1:1). 24h later, the solution was replaced, and the tissue was kept in darkness at room temperature. After two weeks, the tissue was transferred to solution C and stored for 72h. The tissue was then embedded in paraffin, and 200μm sections were obtained in a vibratome, mounted in gelatinized slides and impregnated with solution C. For the staining, the sections were rinsed in Milli-Q water two times for 4 min and transferred to the staining solution (solution D, solution E and Milli-Q water, 1:1:2) for 10 min. The sections were rinsed again in Milli-Q water (2x 4 min), dehydrated (ethanol 50%, 75%, 95%, 4 min each followed by absolute ethanol, 4 times of 4 min) and cleaned (xylene 3 times for 4min each). The slides were covered with Permount mounting medium, and the images were acquired with Visiopharm in an Olympus BX53 and analyzed with IMARIS 8.4.2.

#### 2.3.3 ELISA

Trunk blood was obtained at euthanising and centrifuged (2,000 rpm, 10 min, 4°C) to collect the plasma stored at −80°C. The corticosterone ELISA assay was performed as described by the manufacturer (ENZO Life Sciences). Briefly, the solutions were left 1h at room temperature, whereafter the samples were diluted in sample buffer (1:100), and 100 μl were added to the plate, followed by 50 μl of the blue conjugate. Then, 50 μl of antibody was added, and the plate was incubated at room temperature for 2h in a plate shaker. Following a wash of the plate with 400 μl of washing solution (3 x, 200 μl), pNpp substrate was added to each well, and the plate was incubated for 1h at room temperature. Finally, 50 μl of stop solution was added, and the plate was read at 405 nm using a Clariostar microplate reader (BMG LabTech, Germany).

#### 2.3.4 Western Blot

The HPC was homogenized as previously described (49) and used for dual-fluorescent western blot (50). Briefly, an 8% or 10% NuPAGE Bis-Tris Midi gel (Invitrogen®) in MES or MOPS running buffer (Invitrogen) was used to separate 20 μg of total protein. Later, the separated proteins were transferred to a 0,2 μM nitrocellulose membrane (BioRad trans-blot turbo). The membrane was blocked with Odyssey blocking buffer (OBB) (LI-COR)/TBS (1:1) for 1h at room temperature. Later, the membrane was incubated with the primary antibody diluted in OBB/TBS −0.1% Tween-20 (TBS-T) (1:1) at 4°C overnight; rabbit anti-pTrkB (1:500, Cell Signalling), goat anti-TrkB (1:500, R&D), rabbit anti-pERK1/2 (1:500, Cell Signalling), mouse anti-ERK1/2 (1:1000, Cell Signalling), mouse anti-GR (glucocorticoid receptor, 1:500, Cell Signalling), rabbit anti-PSD95 (post-synaptic density 95, 1:1000, Cell Signalling), mouse anti-actin (1:3000, LI-COR), rabbit anti-BDNF (1:1000, Abcam), rabbit anti-pmTOR (1:500, Cell Signalling), mouse anti-mTOR (1:1000, Cell Signalling), rabbit anti-CREB (1:1000, Cell Signalling) and rabbit anti-FKBP52 (1:1000, Abcam). Then the membrane was washed in TBS-T four times for 5 min and incubated for one hour in the dark at RT with secondary antibodies diluted in OBB/TBS-T(1:1); donkey anti-rabbit 800CW, donkey anti-goat 680RD, goat anti-rabbit 800CW and goat anti-mouse 680RD (LI-COR). The protein levels were assessed using Odyssey CLx Scan (LI-COR).

## 3 RESULTS

### 3.1 Behavioural responses to trauma in FSL, FRL and SD rats

In order to discard potential bias factors in the analysis of memory in the CFC, we evaluated the basal behaviour of the three strains during the habituation to the conditioning box (Figure 1). ANOVA analysis revealed no differences among strains regarding distance traveled [F (2, 21) = 1.629, p=0.2199], mean velocity [F (2, 21) = 1.629, p=0.2200], rearing [F (2, 21) = 2.000, p=0.1603], jumping [F (2, 20) = 1.922, p=0.1724] and basal freezing [F (2, 21) = 0.8202, p=0.4539]. However, there were statistically significant differences in grooming [F (2, 20) = 8.772, p=0.0018]. The pattern of movement was also different among strains during the habituation phase, in which FSL spent more time in the inner zone [F (2, 21) = 9.685, p=0.0010] and less time in the outer zone [F (2, 21) = 9.685, p=0.0010], as revealed by ANOVA followed by Bonferroni post-hoc. This pattern was not observed at the end of the protocol [Inner zone: F (2, 18) = 0.7666, p=0.4792; Outer zone: F (2, 18) = 1.810, p=0.1922]. We also assessed the performance of the three strains in the NOR; as indicated by one sample t-test analysis, both SD [t=3.161, df=13, p=0.0075] and FRL [t=2.241, df=12, p=0.0447] were able to recognize the novel object, this was not observed in FSL [t=0.5080, df=12, p=0.6206].

**Figure 1:**
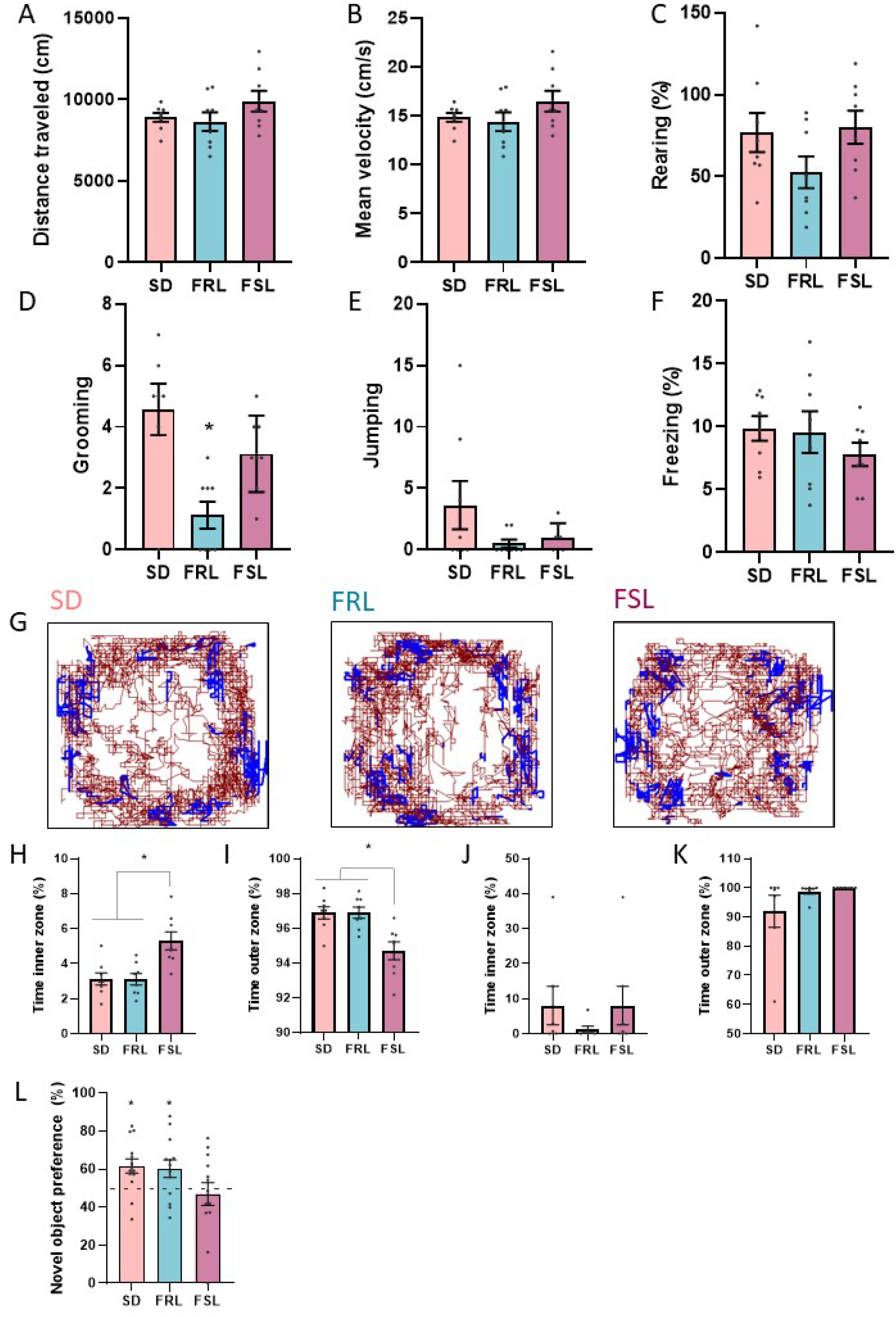
Basal behaviour. A) Distance traveled, n=8-8-8. B) Mean velocity, n= 8-8-8. C) Rearing, n=8-8-8. D) Grooming, n= 7-8-8. E) Jumping, n=8-8-7. F) Freezing, n=8-8-8. G) Representative pattern of movement during the habituation phase for SD, FRL and FSL. H) Percentage of time in the inner zone during habituation n=8-8-8. I) Percentage of time in the outer zone during habituation n=8-8-8. J) Percentage of time in the inner zone during reinstatement test n=7-7-7. K) Percentage of time in the outer zone during reinstatement test n=7-7-7. L) Preference for the novel object, n=14-13-13. * p<0.05

Since differences in performance could derive from differences in shock perception, we evaluated whether the fear response against the shock was different among the strains. The freezing during the conditioning was significantly moderated by time but not by strain [Strain: F (2, 126) = 2.336, p=0.1009; Time: F (5, 126) = 46.40, p<0.0001; Interaction: F (10, 126) = 1.237, p=0.2739] (Figure 2C).

**Figure 2:**
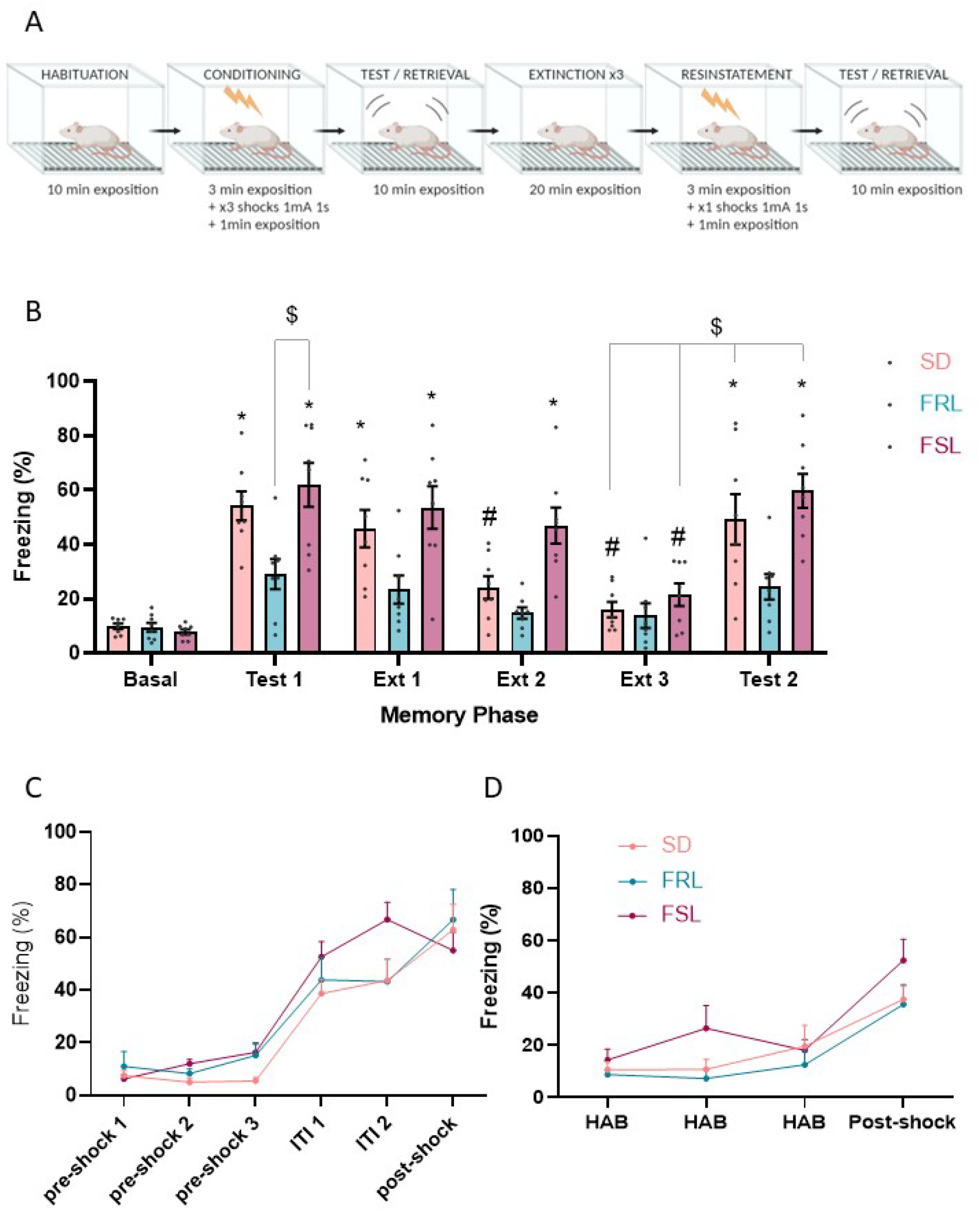
Levels of freezing in the contextual fear conditioning. A) Experimental design. B) Levels of freezing in different phases of the CFC, n=8-8-8. C) Levels of freezing by minute in the conditioning phase of the CFC, n=8-8-8. D) Levels of freezing by minute during the reinstatement phase of the CFC, n=8-8-8. * p<0.05 compared to basal, # p<0.05 compared to Test 1, $ p<0.05 as indicated by the grey lines

We next characterized the performance of the three strains during different memory phases in the CFC (Figure 2A-B). Two-way ANOVA showed significant effect as well as interaction of strain and memory phase [Strain: F (2, 126) = 27.22, p<0.0001; Phase: F (5, 126) = 26.12, p<0.0001; Interaction: F (10, 126) = 2.553, p=0.0077]. Post-hoc analysis revealed that FRL animals presented lower levels of freezing during the test when compared to SD or FSL. Although FSL did not differ from SD during the test, this strain presented impairments in the extinction of fear memory (Figure 2B). During the reinstatement session, freezing was moderated by time point but also by strain [Strain: F (2, 84) = 4.487, p=0.0141; Time: F (3, 84) = 18.12, p<0.0001; Interaction F (6, 84) = 0.6760, p=0.6694] (Figure 2D). Lastly, in the reinstatement test, the freezing levels were similar between FSL and SD, but again FRL presented lower levels of the conditioned response (Figure 2B).

To further evaluate the performance in fear conditioned models, we perform the delay auditory fear conditioning, a task independent of the HPC (51). We observed that during the conditioning FSL presented higher levels of freezing that FRL in response to tone+shock [Strain: F (1, 48) = 6.728, p=0.0125; Time: F (3, 48) = 33.69, p<0.0001; Interaction: F (3, 48) = 0.9064, p=0.4450] (Supplementary Figure 1B). During the tone test, two-way ANOVA revealed significant effect of strain and time [Strain: F (1, 52) = 8.956, p=0.0042; Time: F (3, 52) = 18.61, p<0.0001; Interaction: F (3, 52) = 2.412, p=0.0772] (Supplementary Figure 1C). However, no differences were observed among strains in the levels of freezing during the contextual test [t=1.434, df=12, p=0.1771] (Supplementary Figure 1D).

### 3.2 Differences in hippocampal morphology between FSL, FRL and SD rats

Considering the differences in performance among the strains during CFC, possible alterations in hippocampal neuronal morphology sustaining the behavioural pattern, especially the differences between FRL and FSL-SD, were tested. Representative images from Golgi-cox staining and IMARIS neuronal reconstruction can be seen in Figure 3A-B. When compared to SD, FSL and FRL displayed more dendrites [F (2, 113) = 11.68, p<0.0001] and dendritic spines [F (2, 113) = 5.532, p=0.005], although no differences were found regarding spine density [F (2, 114) = 1.054, p=0.3520]. Following classification, significant differences in the number of spines concerning filopodia [F (2, 115) = 4.408, p=0.0143], long-thin [F (2, 114) = 4.178, p=0.0177] and mushroom [F (2, 115) = 4.053, p=0.0199] types but not stubby [F (2, 112) = 1.862, p=0.1601] were observed. Differences among strains were also observed regarding the mean length of the dendrites [F (2, 115) = 7.815, p=0.0007] (Figure 3J).

**Figure 3:**
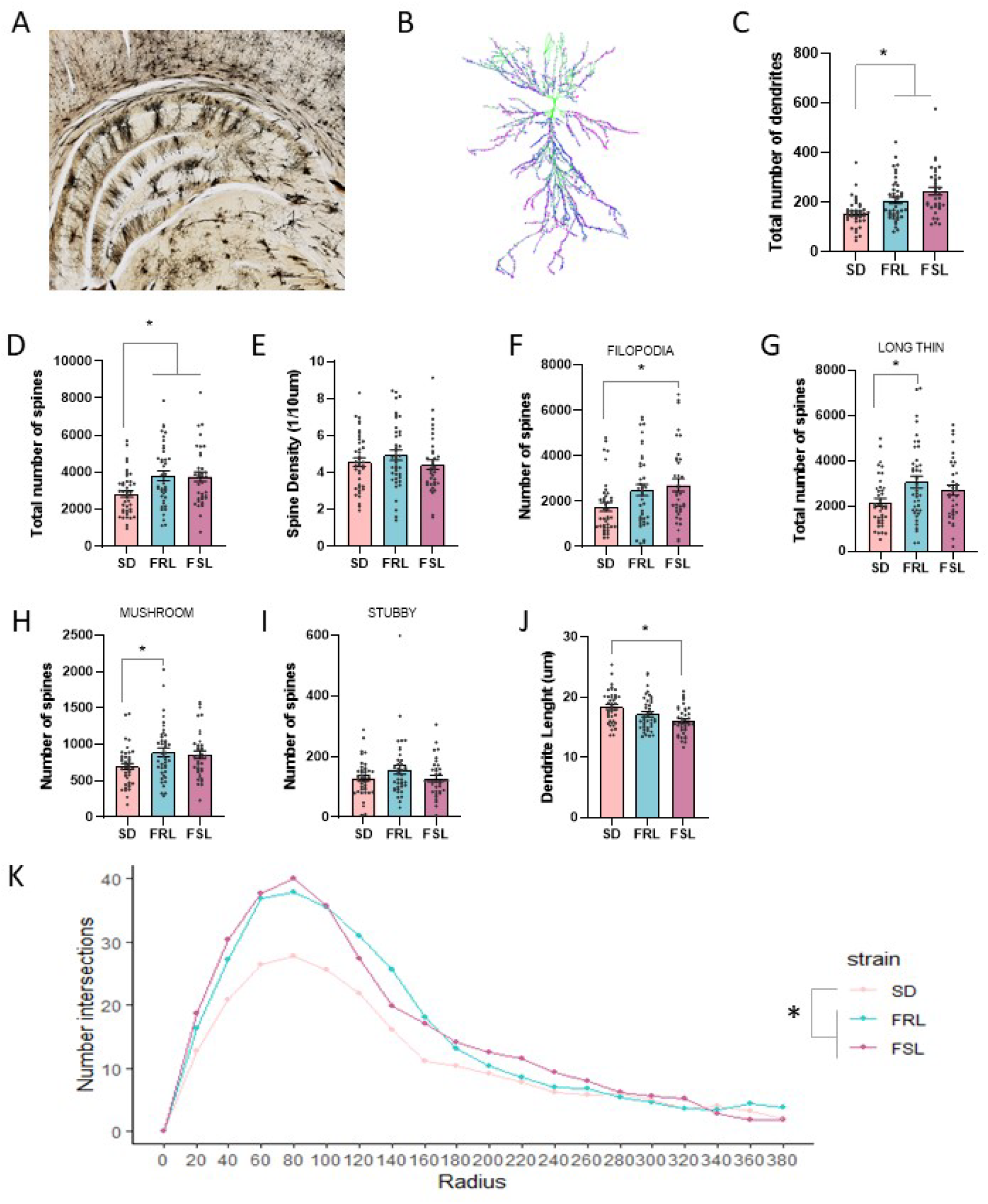
Morphological analysis of dorsal hippocampal CA1 neurons. A) Representative image of Golgi-Cox staining B) Representative image from IMARIS 3D reconstruction. C) Total number of dendrites by neuron, n=39-41-36. D) Total number of spines, n=40-40-36. E) Spine density, n=40-41-36. F) Filopodia number of spines, n=40-41-37. G) Long thin number of spines, n=40-41.36, H) Mushroom number of spines, n=40-41-37. I) Stubby number of spines, n=39-40-36. J) Mean dendrite length, n=40-41-37. K) Number of intersections from SHOLL analysis. p>0.05 as indicated by the grey lines.

**Figure 4:**
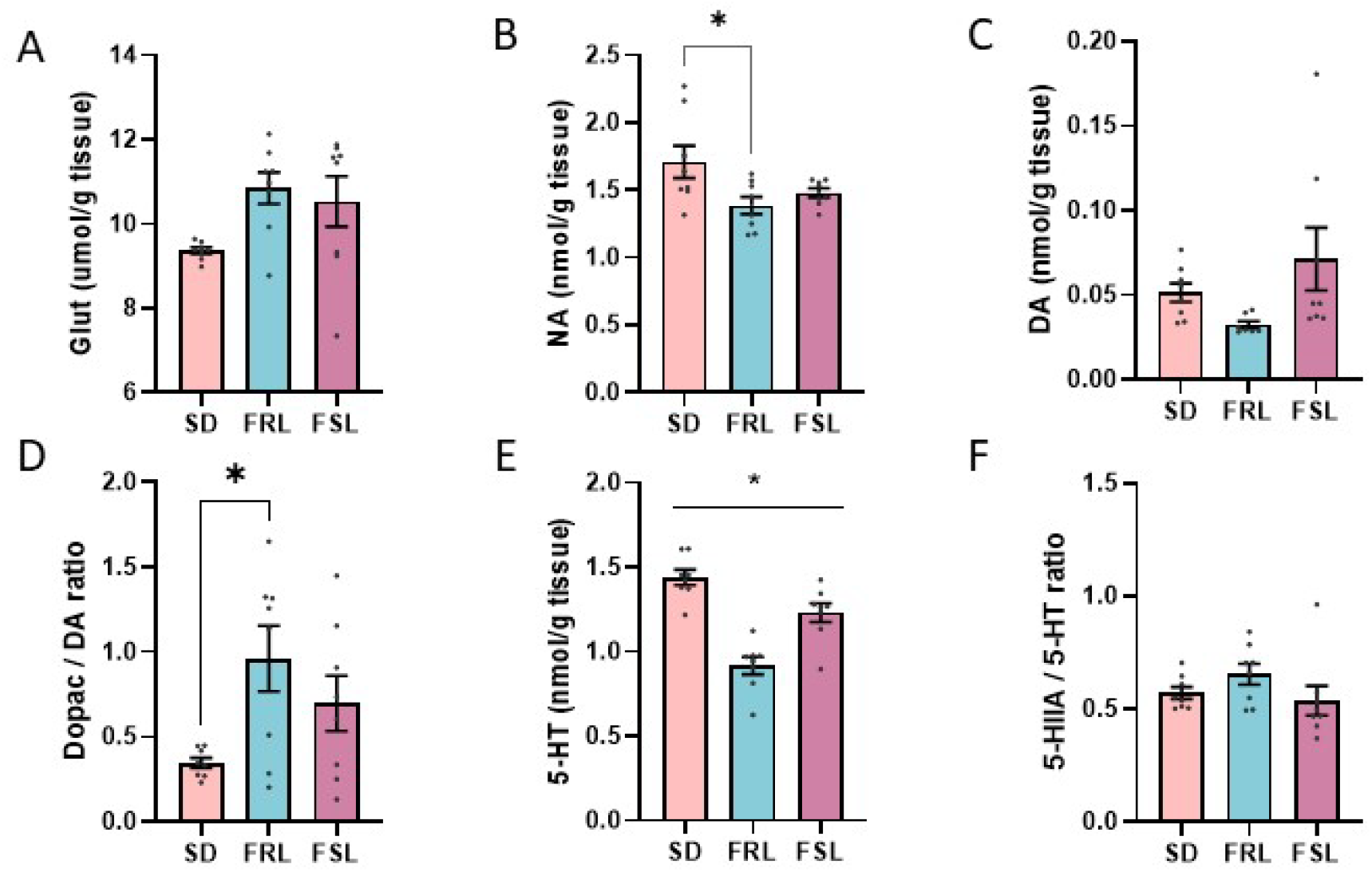
Levels of monoamines in the HPC and PFC. A) Levels of glutamate in the HPC, n=8-8-8. B) Levels of noradrenaline in the HPC, n=8-8-8. C) Levels of dopamine in the HPC, n=8-7-8. D) Dopamine turnover in the HPC, n=8-8-8. E) Levels of serotonin in the HCP, n=8-8-8. F) Serotonin turnover in the HPC, n=8-8-8. p>0.05 as indicated by the grey lines.

Two-way ANOVA of the sholl analysis showed that the number of intersections was moderated by radius, strain and strainXradius [Strain: F(2,2042) = 25.283, p<0.001; Radius F(1,2042) = 554.303, p<0.001; Interaction: F(2,2042) = 8.047, p>0.001], post-hoc analysis suggested that FRL and FSL presented higher complexity of the dendritic tree when compare to SD (Figure 3K).

### 3.3 Neurotransmitter levels in the hippocampus of FSL, FRL and SD rats exposed to trauma

Considering that neurochemical imbalances in the HPC have been associated with CFC (52, 53), we analyzed the levels of neurotransmitters in the HPC of rats submitted to the CFC. In the HPC, significant differences were observed in the levels of glutamate [F (2, 21) = 3.631, p=0.0442], noradrenaline [F (2, 21) = 4.214, p=0.0289] and serotonin [F (2, 21) = 26.42, p<0.0001] but not in the 5-HT turnover [F (2, 21) = 1.567, p=0.2322]. On the contrary, levels of dopamine were not different [F (2, 20) = 2.676, p=0.0934] but significant differences were found in the dopamine turnover [F (2, 21) = 4.407, p=0.0252].

### 3.4 Molecular pathways potentially involved in the FSL performance in the CFC

Since neuronal morphology and neurotransmitter levels after the CFC could not explain the behavioural pattern observed, it was hypothesized that FSL deficits in the extinction and FRL deficits in acquiring and expressing the contextual fear memory could be due to alterations during the consolidation of the original fear-memory. Thus, potential pathways involved with stress response and hippocampal plasticity were evaluated. Two-way ANOVA revealed that CORT levels were moderated by strain and conditioning protocol [Strain: F (2, 26) = 6.562, p=0.0049; Conditioning: F (1, 26) = 4.265, p=0.0490; Interaction: F (2, 26) = 0.2221, p=0.8023], although GR expression was only moderated by strain [Strain: F (2, 18) = 6.566, p=0.0072; Conditioning: F (1, 18) = 0.9988, p=0.3308; Interaction: F (2, 18) = 0.3854, p=0.6856] (Figure 5A), suggesting that higher levels of CORT are present in the group exposed to the shock when compared to the context only group, and that the CORT levels differs among strains (FRL>FSL>SD). Conversely, levels of GR in the HPC were drastically lower in FRL when compared to FSL or SD (Figure 5B). Thus, in light of these results, the expression of FKBP51, an indicator of GR functioning (54), was evaluated. Two-way ANOVA revealed a significant effect of strain but not conditioning protocol or interaction between factors [Strain: F (2, 18) = 8.340, p=0.0027; Conditioning: F (1, 18) = 0.3360, p=0.5693; Interaction: F (2, 18) = 0.1412, p=0.8692] (Figure 5C).

**Figure 5:**
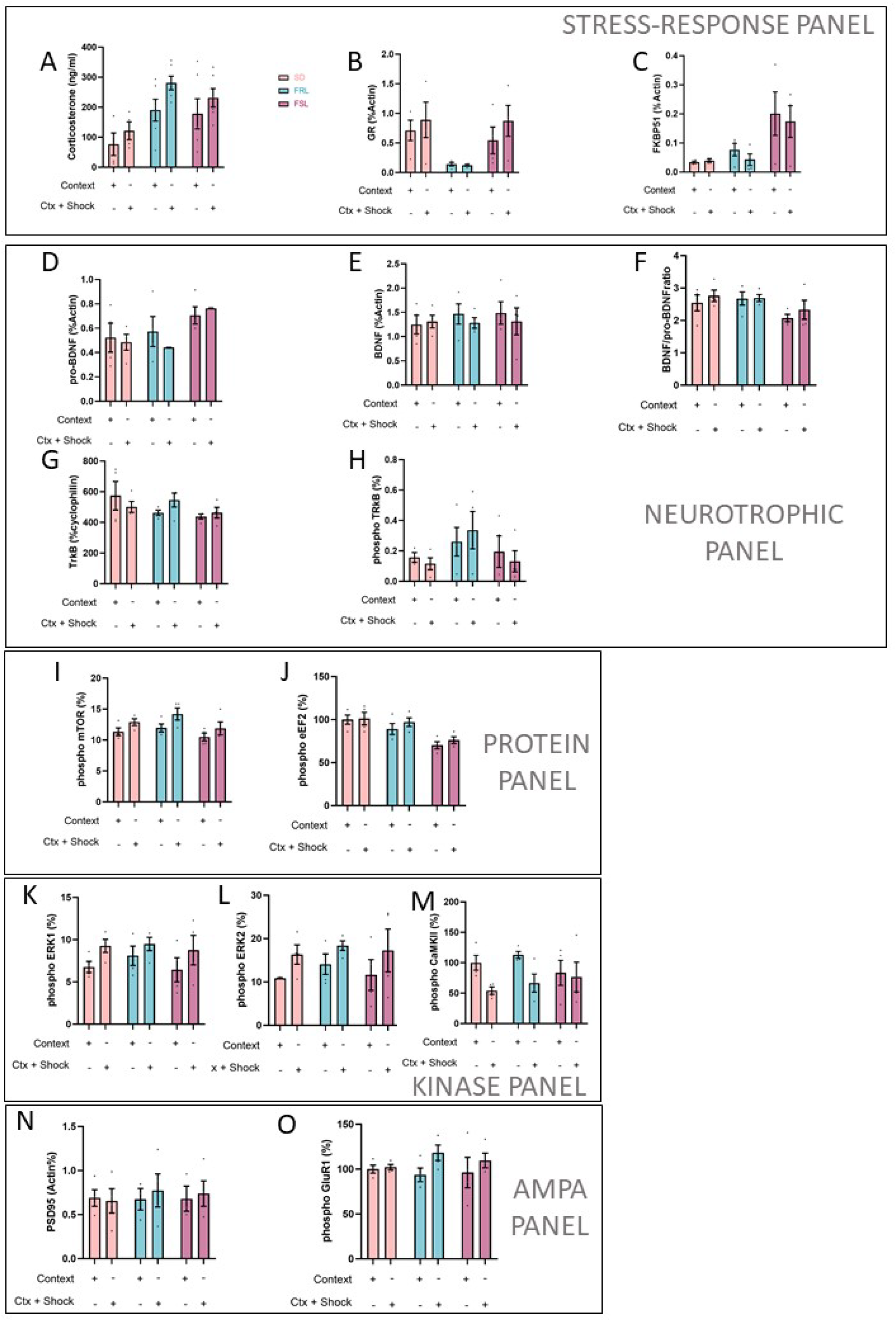
Molecular analysis. A) Corticosterone levels in serum, n=4-4-6-6-6-6. B) Glucocorticoid receptor expression, n=4. C) Total FKBP51, n=4. D) Total pro-BDNF, n=4-4-4-3-4-3. E) Total BDNF, n=4. F) BDNF/proBDNF ratio, n=4. G) Total TrkB, n=4. H) Phospho-TrkB, n=3-4-4-4-4-4. I) Phospho-mTOR, n=4. J) Phospho-eEF2, n=4. K) Phospho-ERK1, n=4. L) Phospho-ERK2, n=3-4-4-4-4-4. M)Phoshpo-CaMKII, n=4. N) Total PSD95, n=4. O) Phospho-GluR1, n=4. *p<0.05 as indicated by the grey lines.

Finally, plasticity pathways modulated by CORT and GR functioning were analyzed to understand if the endocrine differences between the strains could result in molecular changes associated with stress response and memory processing. First, we evaluated neurotrophin signaling: BDNF [Strain: F (2, 18) = 0.2033; p=0.8179; Conditioning: F (1, 18) = 0.3671, p=0.5521; Interaction: F (2, 18) = 0.2383, p=0.7904], pro-BDNF [Strain: F (2, 16) = 4.252, p=0.0330; Conditioning: F (1, 16) = 0.2584, p=0.6181; Interaction: F (2, 16) = 0.5234, p=0.6023], BDNF/pro-BDNF ratio [Strain F (2, 18) = 3.573, p=0.0493; Conditioning: F (1, 18) = 0.9576, p=0.3408; Interaction F (2, 18) = 0.1995, p=0.8209] (Figure 5). Then, we analyzed kinases extensively associated with consolidation: pERK1/2 [Strain: F (2, 18) = 0.5977, p=0.5606; Conditioning: F (1, 18) = 5.840, p=0.0265; Interaction: F (2, 18) = 0.08700, p=0.9171] and pCaMKII [Strain: F (2, 18) = 0.3700, p=0.6958; Conditioning: F (1, 18) = 6.769, p=0.0180; Interaction: F (2, 18) = 1.064, p=0.3659]. Since CaMKII is associated with AMPA receptor trafficking, which is central for synaptic plasticity and memory (55), we evaluated phosphorylation of GluR1; pGluR1 [Strain: F (2, 18) = 0.1376, p=0.8724; Conditioning F (1, 18) = 3.166, p=0.0921; Interaction: F (2, 18) = 0.7248, p=0.4980] and PSD95 [Strain: F (2, 16) = 0.07021, p=0.9325; Conditioning: F (1, 16) = 0.1228, p=0.7306; Interaction: F (2, 16) = 0.1139, p=0.8931]. Finally, we assessed the phosphorylation of mTOR-eEF2 as an indicative of increased protein synthesis: pmTOR [Strain: F (2, 18) = 3.031, p=0.0734; Conditioning: F (1, 18) = 7.583, p=0.0131; Interaction: F (2, 18) = 0.1807, p=0.8362] and eEF2 [Strain: F (2, 18) = 13.34, p=0.0003; Conditioning: F (1, 18) = 1.277, p=0.2734; Interaction: F (2, 18) = 0.2026, p=0.8184].

## 4 DISCUSSION

The present study characterized the behaviour of FSL and FRL in the CFC and the molecular pathways potentially involved in their aberrant behavioural performance. The most important findings were: i) although FSL and SD had a similar conditioned response in the test, FSL required more extinction sessions to significantly reduce the levels of freezing; ii) FRL were unable to display proper levels of freezing during memory retrieval even with an intact innate fear response and contextual memory; iii) Both FSL and FRL present cortical and hippocampal alterations in monoamine and glutamate levels together with an increase in the total number of spines in CA1 when compared to SD; iv) There were no significant alterations in critical plasticity pathways among the strains, but substantial differences in the HPA axis response and glucocorticoid activity in the HPC: FRL animals released higher levels of corticosterone than FSL, along with a drastic decrease in GR expression in the HPC; however, FKBP51 levels were drastically incremented in FSL when compared to FRL or SD. Importantly, we observed that the three strains tested (SD, FRL and FSL) had no confounding baseline differences, such as exploratory behaviour or the innate response to a shock, except for FSL spending more time in the centre of the field when compared with the other strains. This indicates that the behavioural differences between the strains could not result from baseline characteristics.

In contrast with some previous reports (56), but in agreement with others (57), we did not observe a marked hypolocomotion in FSL; in fact, most of the basal behaviours evaluated were not significantly different among strains. In addition, FSL animals also presented an increase in the time spent in the centre of the box compared to the control strains, a trait often associated with low anxiety-like levels (58). Prior studies addressing innate anxiety in FSL reported contrasting results, e.g. no differences or lower anxiety levels when compared to FRL (36, 59, 60) and higher or lower anxiety compared to SD (61). We speculate that the conflicting results can be attributable to varying experimental conditions.

We report for the first time that FRL performs similarly to SD in the object recognition test, indicating a similar recognition index and preserved cognitive skills. Our data also extend prior findings suggesting poor performance of FSL in low-emotional load memory tasks(36–41). In conditioned fear responses, only limited data is available regarding the FSL and FRL rats. In operant conditioning models, FSL showed impairments in the active (62) and passive avoidance tasks (63) compared with FRL. This contrasts with the current findings, where we in the CFC observed lower conditioned responses in FRL and similar levels of freezing in FSL compared to SD. Methodological details may be responsible for this discrepancy. For example, it is possible that the low intensity of the conditioning training (0.4 – 0.8mA) used in the previous studies may not be enough to overcome the well-documented cognitive deficits of FSL (36–41), and/or to trigger resilience mechanisms in FRL. Moreover, active avoidance tests investigate active coping strategies that may differ from passive strategies and unescapable situations, as described in the earlier literature. However, further studies are required to confirm these suggestions.

Interestingly, extinction impairments in FSL were observed when compared to SD. Such impairments are hallmark features in PTSD patients and valid animal models of PTSD (29, 30, 64). To further elucidate the performance of classical fear conditioning tasks, we evaluated the behaviour of FSL and FRL in the delayed auditory fear conditioning, a task unrelated to hippocampal functioning (51). The observation was that the freezing levels of FSL during the test were higher than FRL and that FSL displayed increased freezing after tone+shock compared to FRL, a difference not found when the strains were presented only to the shock+context. These findings suggest that despite the FRL not presenting deficits in other HPC-dependent memories, they could not display appropriate conditioned responses in fear-memory tests. In contrast, FSL animals displayed poor performance in HPC-dependent memory tasks and properly conditioned responses but impairments in the acquisition of safe-contextual learning. The mechanisms involved in these abnormalities seem to be linked to dysfunctions in the HPA axis function. Indeed, HPA axis hypofunction is often described in PTSD patients (65). However, in response to trauma, reminders or threats, PTSD patients present an exacerbated release of cortisol, which can reflect a defective inhibitory control of the HPA axis (66). Cortisol/corticosterone increases the consolidation of contextual memories in U-inverted shape (67, 68). Thus optimal corticosterone levels and HPA functioning are crucial for memory and cognition (69, 70). The low basal cortisol levels in PTSD patients have been suggested as an underlying part of the cognitive deficits in these subjects (71).

Previous studies suggest that FSL animals are also characterized by HPA hypofunction (72, 73). We observed that FSL animals, despite their poor performance in HPC-dependent memory tasks, such as the NOR, presented intact acquisition and retrieval of memories related to fearful stimulus, which can be related to the sub-optimal activity of the HPA axis during non-emotional tasks. In contrast, compared to SD, they display an exacerbated release of corticosterone during the consolidation of traumatic-like memories, with higher levels of FKBP51. Other models of PTSD, such as single prolonged stress (SPS), seem to induce some behavioural effects through GR. For example, SPS-enhancement of fear conditioning or LTP impairment was prevented by GR antagonism prior to stress (74). Moreover, fear conditioning has been shown to enhance the levels of FKBP51 in a sustained way,, thereby decreasing FKBP51 in the IL impaired consolidation and enhanced fear extinction (75). Similarly, the downregulation of FKBP51 in the HPC seems to mediate resilience against specific insults (76, 77). Importantly, FKBP51 seems not to be involved in other HPC-dependent memories (78) and this protein seems to participate in antidepressants’ effect (79, 80). Similarly, serotonin reuptake inhibitors, main pharmacological choice for the treatment of PTSD and MDD, seems to act rescuing some of these hippocampal impairments (25, 81–83) and this was already related with the modulation of GR (25, 84). On the other hand, FRL animals, despite the high levels of corticosterone in plasma showed a drastically low expression of GR in the HPC and similar levels of FKBP51 than SD. Since CORT presents an U-inverted shape in memory consolidation, the high levels of CORT found in FRL combined with low expression of GR/FKBP51 may explain the absence of conditioned responses in these animals. More importantly, differently from FSL, FRL animals presented intact cognitive function, thus the low response cannot be associate to HPC deficits in encoding or retrieved contextual memories.

Finally, our protocol enhanced the phosphorylation of several targets, but we could not find differences among strains. This agrees with previous reports describing no differences in BDNF, phosphorylation of TrkB, CaMKII or PSD95 levels in flinders (63). However, other authors suggested lower hippocampal BDNF basal levels in FSL compared to FRL (85).

Our study presents some limitations that should be taken into consideration. One important aspect is that our experiments did not establish a causal relationship between the HPA-HPC alterations and the behavioural phenotype. Furthermore, our experimental sample did not represent females since we only used adult male rats. Further experiments are, thus, necessary to clarify these issues.

In conclusion, the flinders (FSL and FRL) are valuable tools to study the neural substrates involved in susceptibility and resilience to developing behavioural consequences of trauma exposure. Furthermore, our initial findings indicate that the HPA-HPC function may be an essential system regulating vulnerability and resilience against traumatic events

---

**Supplementary figure 1:**
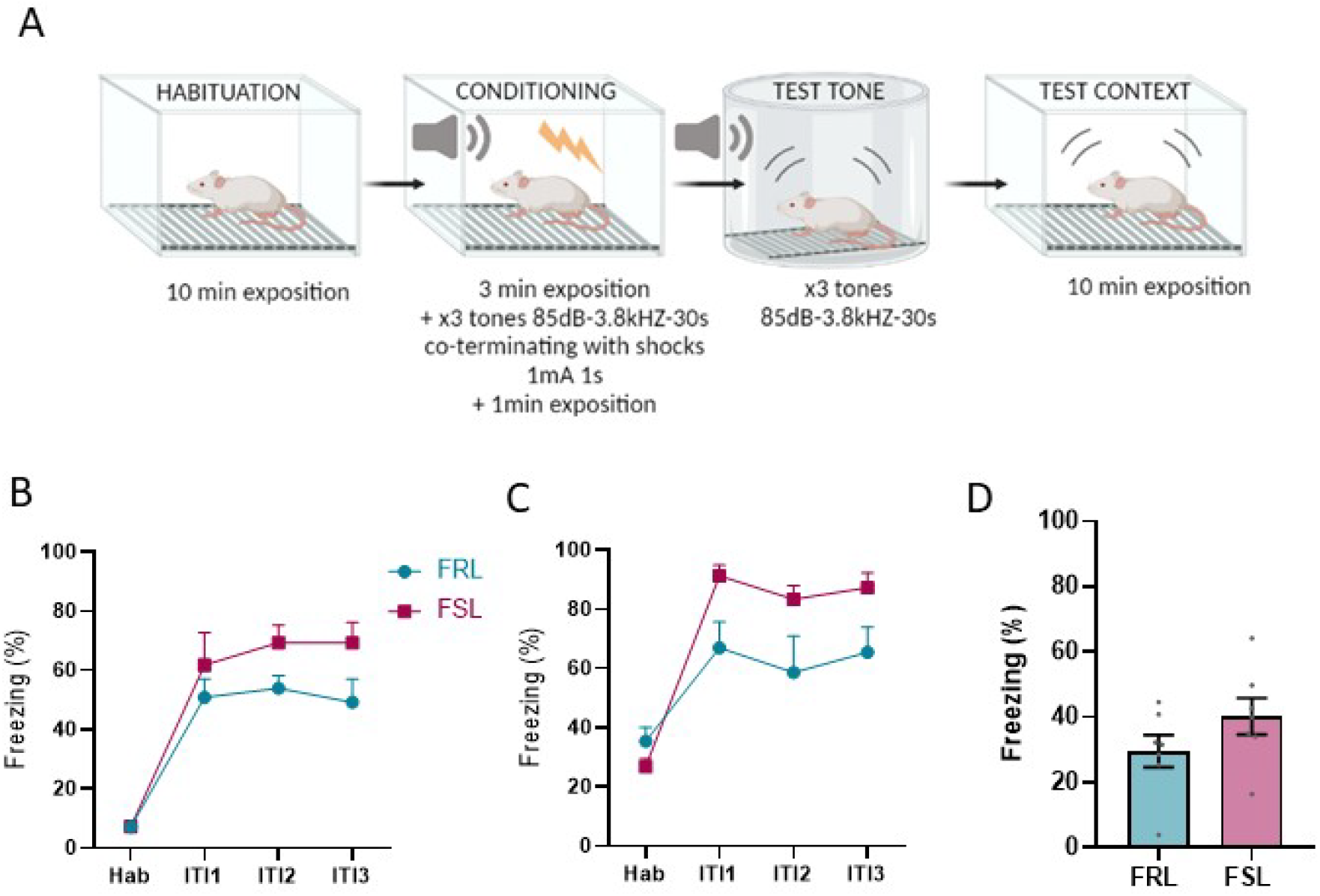
Further characterization of flinders memory performance, results from delay AFC. A) Experimental design AFC. B) Freezing levels during the conditioning phase, n= 7. C) Freezing levels during retrieval of memory in response to the tone, n=8-7. D) Freezing levels during retrieval of memory in response to the context, n=7.

## Notes

### Competing Interest Statement

The authors have declared no competing interest.

